# The eyes have it: Inter-subject correlations of pupillary responses for audience response measurement in VR

**DOI:** 10.1101/2024.01.22.576685

**Authors:** Ralf Schmälzle, Juncheng Wu, Sue Lim, Gary Bente

**Affiliations:** Department of Communication, Michigan State University

**Keywords:** reception analysis, audience response measurement, pupillometry, intersubject correlation, media exposure, media psychology, psychophysiology, virtual reality

## Abstract

The eye is the vanguard of the reception process, constituting the point where visual information arrives and is transformed into neural signals. While we view dynamic media contents, a fine-tuned interplay of mechanisms causes our pupils to dilate and constrict over time - and putatively similarly across audience members exposed to the same messages. Research that once pioneered pupillometry did actually use dynamic media as stimuli, but this trend then stalled, and pupillometry remained underdeveloped in the study of naturalistic media stimuli. Here, we introduce a VR-based approach to capture audience members’ pupillary responses during media consumption and suggest an innovative analytic framework. Specifically, we expose audiences to a set of 30 different video messages and compute the cross-receiver similarity of pupillometric responses. Based on this data, we identify the specific video an individual is watching. Our results show that this ‘pupil-pulse-tracking’ enables highly accurate decoding of video identity. Moreover, we demonstrate that the decoding is relatively robust to manipulations of video size and distractor presence. Finally, we examine the relationship between pupillary responses and subsequent memory. Theoretical implications for objectively quantifying exposure and states of audience engagement are discussed. Practically, we anticipate that this pupillary audience response measurement approach could find application in media measurement across contexts, ranging from traditional screen-based media (commercials, movies) to social media (e.g., TikTok and YouTube), and to next-generation virtual media environments (e.g., Metaverse, gaming).

## Introduction

The eye is the vanguard of the reception process, the point where visual media content arrives at the senses and gets converted into neural impulses. Accordingly, studies of media reception mechanisms leverage eye-tracking to capture the eyes’ response to on-screen information. The current study uses pupillometry to track exposure to media messages, focusing on whether constrictions and dilations of viewers’ pupils can serve as an index to decipher the message to which they are exposed.

The paper is organized as follows: First, we review the functional role and neurocognitive mechanisms of the pupil in the reception process. Next, we introduce general measurement principles and the recent trend to include pupillometric measurement capabilities, particularly in VR headsets. Then, we discuss past applications and current uses of pupillometry in media psychology, vision research, and cognitive science more broadly. Finally, we introduce the current study in which people are viewing a variety of video messages on a (virtual) screen while pupillometric measures are taken, followed by methods, results, and discussion.

### Pupil Dynamics: Function, Mechanism, and Measurement

The pupil - the black, circular opening at the center of the iris - is an essential component of the human eye and gatekeeper for the visual system. Its primary function is to regulate the amount of light entering the eye (Laeng & Alnaes, 2019). Specifically, when the level of light is high, the pupils constrict to limit the amount of incoming light; when the level of light is low, they dilate to absorb as much light as possible (pupillary light reflex; PLR; Ellis, 1981). In addition to lighting conditions, pupil size is sensitive to variations in arousal (Hess & Polt, 1960; Mathôt, 2018; Nunnally et al., 1967; Peavler & McLaughlin, 1967). The muscles of the iris control the size of the pupil and the underlying neural circuitry of these systems is increasingly known (Clewett et al., 2018; Sirois & Brisson, 2014). Thus, when we process a video, pupils change over time in a way that reflects the connections between visual perception and cognition.

Pupillometry measures how pupil diameter changes over time (Mathôt, 2018). Pupillometry has been used for basic and applied purposes, such as neurological research on reflexes, alertness, and workload monitoring (Laeng & Alnaes, 2019; Sirois & Brisson, 2014). Methodological advantages include that the measure is temporal, providing a moment-to-moment readout of reactions to an unfolding stimulus, such as a continuous video. Second, pupillometry is unobtrusive in that recording is passive; thus, it requires no interruption of ongoing reception processes, which could introduce confounds. Third, as a nonverbal measure, pupillometry circumvents potential response biases and other difficulties with introspection. These advantages suggest pupillometry as an attractive measure to examine how people respond to media.

Pupillometry has been previously suggested for media research but remains underdeveloped. The pioneering work of E. Hess constitutes perhaps the first and most well-known applicationsof pupillometry to media studies. In one of many studies, Hess measured viewers’ pupils while they watched a Western series (de Winter et al., 2021; Hess, 1975). Although Hess’ work helped popularize pupillometry, most subsequent applications lie in cognitive psychology (Mathôt, 2018) rather than media psychology. For example, pupillometry played a role in seminal work on cognitive load and effort measurement (Beatty, 1982; Kahneman & Beatty, 1966), which is relevant for media research, and the method gets used in neighboring fields (Laeng et al., 2016; Piquado et al., 2010). However, it appears that the application of pupillometry to examine responses to media has stalled and no work has been published in the Journal of Media Psychology in the last decades.

### Pupillometry’s Untapped Potential for Media Response Measurement

Recent trends in eye-tracking technology suggest pupillometry as a promising parameter for media response measurement. As of 2023, technology has matured to a degree where the once bulky lab equipment is commodified and even integrated into wearable devices. For instance, mobile eye-tracking glasses enable researchers to study pupillary changes during video consumption (Ehinger et al., 2019; Ferhat & Vilariño, 2016; Steil, 2019). Moreover, there is a strong trend to integrate eye-tracking into VR headsets, which makes the method far more available, affordable, and scalable. Coupled with an expected trend toward VR-based advertising, we can thus expect an uptick in interest in eye-tracking for media response measurement. This will implicitly promote pupillometry because it relies on the same technology as coordinate-based eye-tracking.

But aside from availability and desirable measurement properties, the central question is what theoretical benefits we can gain from pupillometry? Our answer is simple: We argue that by capitalizing on pupillometry’s most well-established and robust phenomenon - the pupillary light reflex (PLR) - it is possible to turn pupillometry into a method for audience response measurement. Doing so could have implications for objectively quantifying exposure and states of audience engagement, which are key constructs in media and communication research (Biocca et al., 1994; de Vreese & Neijens, 2016). This argument is laid out next.

When the level of illumination increases, the pupil constricts. Conversely, when the level of illumination decreases, the pupil dilates. This relationship is governed by the pupillary light reflex (PLR), an automatic response to changes in lighting conditions that regulates visual sensitivity. In most psychology-focused applications, this pupillary light reflex tends to be treated as a confound^1^ (Mathôt, 2018). By contrast, our proposal is that we can directly mine the full pupillometric information - including the PLR-based responses. Specifically, we argue that depending on the goal of the research, one can treat the PLR responses as *signal* instead of *noise*. The rationale behind this reasoning is as follows: A video message consists of time-varying information. When this information arrives at the eye, it will evoke dilations and constrictions of the pupil over time. Critically, the general nature of these mechanisms suggests that when different members of an audience process the same video, their pupils should constrict and dilate similarly over time, revealing correlated fluctuations across viewers. In the context of human neuroimaging, this principle is sometimes labeled as audience coupling or brain synchronization; the underlying approach is known as inter-subject correlation analysis (Hasson et al., 2010; Schmälzle & Grall, 2020; Wang et al., 2012). In a nutshell, our proposal is to adopt this approach for the analysis of pupillometric data. Of note, such pupil-ISC has already been demonstrated to be generally feasible, but the theoretical connections for media response measurement have not been articulated, and it is unclear whether and to which degree the pupillary responses can be used to decode which video a person is viewing (Golland et al., 2014; Madsen et al., 2021; Madsen & Parra, 2022).

To summarize, we propose that by comparing pupillometric data across viewers, one could objectively track exposure to specific messages during the ongoing reception process. This has been a goal of media measurement for decades (Biocca et al., 1994; de Vreese & Neijens, 2016). Achieving this goal could thus have important theoretical consequences for quantifying the nexus between exposure and reception processes and practical potential for audience response measurement.

### The Current Study and Hypotheses

Visual media change over time, which in turn triggers changes in pupil dilation. When multiple viewers comprising an audience are exposed to the same video messages, we can, therefore, expect that their pupils should fluctuate similarly (see Figure 1).

**Figure 1.**
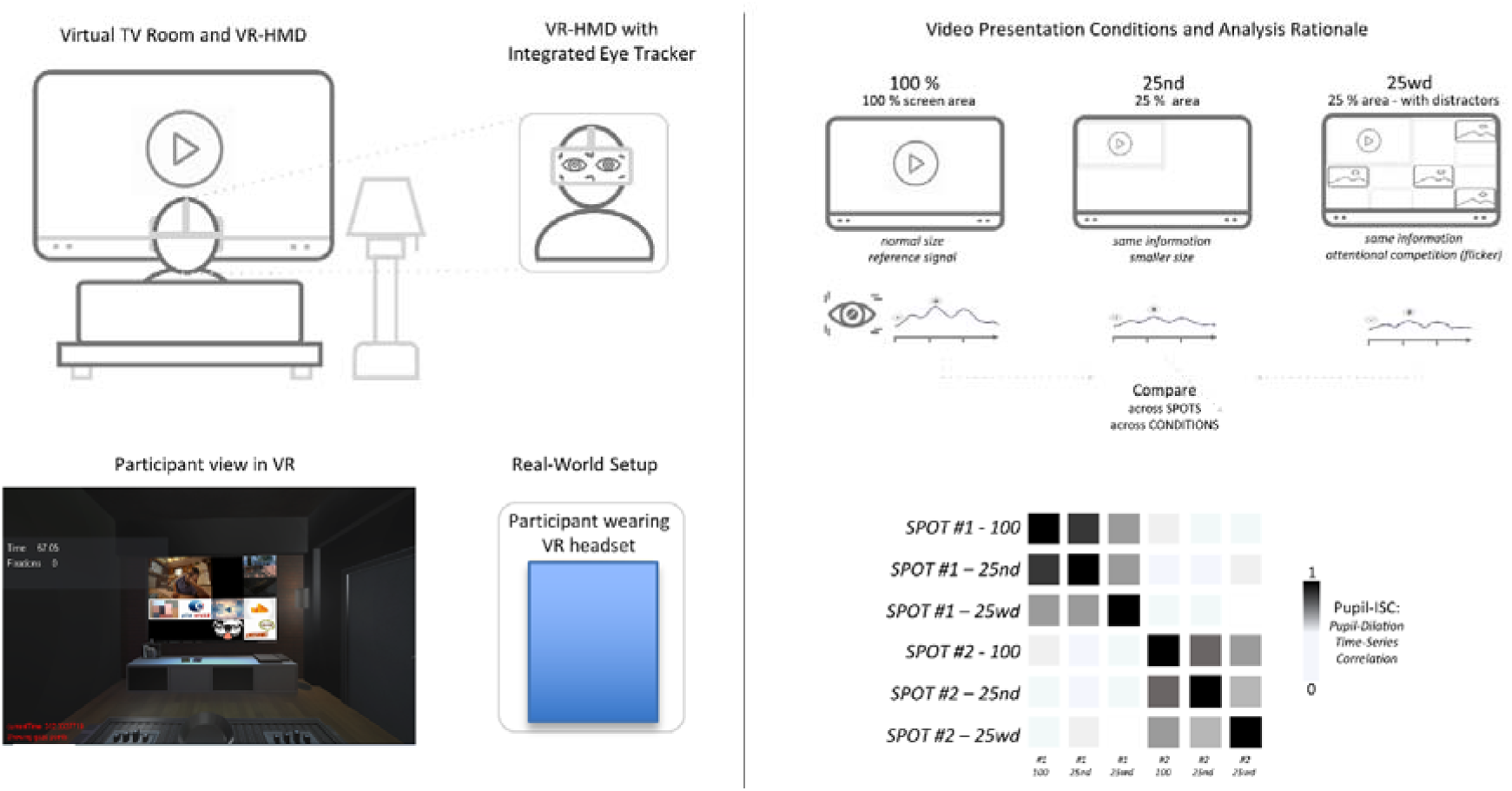
Study Overview. Top left panel: VR head-mounted device (HP Omnicept) affords the seamless delivery of the experimental video stimuli with integrated measurement potential. The device has a high-quality eye tracker that captures pupil dilation at 50 Hz. Top right panel: In the VR environment, a TV/living room setup can be realized, and the experimental videos are presented on a wall-mounted monitor inside VR - just like in a real living room. This setup controls the environment, making it identical for every participant, but preserves the realism of the TV viewing situation. Top right panels: 30 spots of commercials and health-related PSAs are presented in three different conditions: Either as a full-screen video (100%), as a shrunk version of the video (25%nd), or as a shrunk version of the video surrounded by flickering distractor images (25%wd), mimicking the situation in which banner ads would pop up around the video. With this setup, we can compare the similarity of pupillary response traces across the different video conditions (because video presentations are counterbalanced across participants) as well as across different videos. Bottom right panel: The rationale of using Pupil-ISC analysis is that viewing the same spot in different conditions should elicit similar temporal pupil-dilation/constriction profiles, which could be used to decode which spot a participant is seeing.

Based on this reasoning, we studied how the pupils of an audience comprising 60 viewers respond to a collection of video messages. We used a VR-integrated eye tracker, which allowed for simultaneous stimulus delivery (showing videos) and response measurement (capturing pupil dilation). Notably, VR offers another benefit that is worth highlighting: to manipulate and control the viewing conditions. Specifically, we used VR to create a virtual media reception lab comprising a stylish TV viewing room with a (virtual) wall-mounted screen on which the videos were displayed.

Our overarching hypothesis was that pupillometry traces during exposure to specific videos would correlate across viewers. In turn, mining this signal should enable above-chance decoding of a given video’s identity. We thus predicted that by collecting a repertoire of pupillometry data from other users exposed to several videos, one could predict which video a new user is viewing. If true, then this would have implications for the measurement of exposure, a foundational construct for mass communication and a key variable in advertising (de Vreese & Neijens, 2016; Hornik, 2002; Smit & Neijens, 2011).

To further examine the boundary conditions of this expected effect, we manipulated the videos in different conditions (between subjects): First, the video was presented at normal size, filling the entire screen (100%). Two additional conditions presented a shrunk version of the same videos (25%, like, e.g., a YouTube or TikTok video that appears in the top right corner of the screen), thus retaining the content but shifting overall size and brightness parameters. One condition presented this 25% version of the video without distractors, whereas in another condition, the 25% video was flanked by distracting thumbnail images (the thumbnails appeared and disappeared over the duration of the video, much like in banner advertising on websites). We expected that the pupillary response similarity (and thus decoding performance) should be highest for the original videos (100%), followed by the distraction-free, shrunk version (25%, no distractors), and, finally, the version with distractors (25%, with distractors). To further mimic incidental media viewing conditions, we introduced an unannounced memory test. This way, we can also link the experimental manipulation and the reception-response data to outcomes, particularly to subsequent memory.

## Methods

The data and materials for this study are available at https://anonymous.4open.science/r/vr_video_pupil_study-0CEE/

### Participants

Sixty participants were recruited at a large university in the United States (*mean age* = 20.72, 31 self-identified women). Data for one participant had to be excluded.^2^ All participants provided written informed consent to the IRB-approved protocol and they received course.

### Stimuli, Virtual Viewing Room, and Experimental Procedures

#### Stimuli

30 videos (all 30 seconds long) served as experimental stimuli. 18 video clips had commercial content, 12 were health-related public service announcements. All videos were presented in three types of format: 100% full-screen size, 25% screen size with no distractors, and 25% screen size with distractors. In the 100% condition, the video was shown, filling the entire TV screen. In the 25% condition, the same video was shown on the TV but shrunk in size to fill only 25% of the area. In the 25%-plus-distractors condition, the same video was shown (shrunk to 25%), but this time with added distractor images (see Figure 1 for an illustration). These distractors included brand logos and popular imagery. Distractors were shown to the right and below the actual videos - just like it is common on popular websites like YouTube, Facebook, or Baidu. Each participant viewed 10 videos from each of these three conditions. Messages were selected and presented in a pseudo-randomized order so that across the 60 participants, each video version was viewed by 20 viewers. Every viewer saw the same 30 messages, but each source message was seen in only one of the three format options.

#### VR Viewing Room

We developed a virtual viewing room with a wall-mounted TV to show the videos. The core room model was downloaded from Sketchfab.com and edited in Vizard’s Inspector software. The VR room looked like a real and natural living room with a flat-screen TV on the wall. Participants could walk and take place in a seat on a (virtual) couch (in real life, they were seated on a comfortable chair in the laboratory), right across the wall on which the video started playing. This resembles the increasingly common VR-viewing setups, which feature similar living-room style settings and virtual displays. In a validation study (redacted_for_review), it was established that viewing videos in this virtual environment as compared to a real-life TV viewing setup elicited highly similar psychophysiological and subjective response patterns across audiences.

#### Experimental Procedure

Participants completed a vision test before the VR-integrated eye-tracking was put on and calibrated. For a subsample of participants, we also measured EEG, but these data are independent of the current study and will be reported elsewhere.

The experiment involved passively viewing the 30 videos on the virtual TV screen. Between each video, there was a 9-second short break video featuring a 3-2-1-countdown and a fixation cross to focus participants’ attention. After completing the virtual TV viewing session, which took about 20 minutes, a structured interview asked which videos they remembered (free recall). Finally, participants responded to a brief survey about demographics and their VR experience. In addition to the free-recall, we measured participants’ memory using the frame-recognition method (Rossiter et al., 2001), presenting screenshots of all videos and distractors. Finally, we asked participants survey questions regarding spatial presence (Hartmann et al., 2016), immersive tendencies (Witmer & Singer, 1998), and the occurrence of symptoms while in VR (Kim et al., 2018).

## Measurement and Analysis Methods

### Equipment for VR and Pupillometry

Participants viewed the videos wearing an HP-Reverb-Omnicept-Pro. This HMD contains an integrated Tobii eye-tracker (120 Hz). We used the Vizard software (WorldViz Inc. Santa Barbara) to create the VR environment, develop control procedures, and measure pupil dilation. Pupil dilation data were recorded continuously during the whole VR-viewing-experience, digital triggers marked the on-and off-set of each video, and were saved together with the pupil measurements to a data file.

### Pupillometry Recording and Analysis

The main dependent variable was participants’ pupil dilation over time. Pupil dilation values for each participant, video, and sub-condition were parsed from the resulting data file using Python-based routines, and pupil values missing due to eye blinks were coded as NaNs and subsequently interpolated. All analyses were run in Jupyter notebooks using Python 3.7.

After recording, the data file from each participant was read in, video-related pupil-time-series were extracted, eventual blinks were interpolated, and the data were downsampled to 20 Hz and stored individually for each participant, each of the 30 videos, and each of the three conditions (100, 25nd, 25wd).

To examine the similarity of responses to the same videos and across sub-conditions, we computed ISC analysis between viewers’ pupil dilation data. Specifically, ISC analysis was conducted for each video under the 100% condition and then for the 25nd and 25wd conditions (e.g., all pupillary time series for the 20 participants who viewed the 100% version of the 30s “Milk”-commercial, etc.). ISC analysis was conducted in line with guidelines for neuroimaging measures, which were adapted to the context of pupil measurement (Nastase et al., 2019). We visualize the pupil-timeseries and their correlations using the split-half methods, but under the hood, we carried out also pairwise and leave-one-out ISC analysis (the online code repository documents all analyses in a reproducible manner).

Moreover, we used the captured pupillary data to train a machine-learning model and tested its performance in predicting which of the videos a held-out participant was viewing (i.e., testing whether pupillary data enabled spot decoding).

Lastly, we also explored potential subsequent memory effects by comparing the pupillary responses for videos that were subsequently recalled (or recognized) against those for videos that were forgotten.

### Measurement of Message Recall, Recognition, and Survey Responses

After participants finished the VR-video-viewing session, the headset was removed and participants completed a verbal interview in which they were asked to freely recall as many videos as possible. Finally, they completed a recognition test in which screenshots of the videos were shown along with distractor screenshots. Finally, they answered survey questions regarding presence, immersion, and potential intereference due to VR-related symptoms (see above).

## Results

After the virtual TV viewing session, participants provided free verbal comments and survey-based answers about their experience in VR. In general, they enjoyed the study and described it as realistic, engaging, and immersive. Analysis of survey responses reveals that participants felt spatially present in the virtual room (*mean*_*spatial presence*_ *= 3*.*58, s*.*d. = 0*.*76*, scales range from 1-5) and experienced very little to no discomfort symptoms (*mean*_*discomfort*_ *= 1*.*38, s*.*d. = 0*.*21*). Together, these results confirm that the virtual TV room provided an authentic viewing experience, with the added benefit that their pupillary responses could be captured throughout exposure and without interference.

### Strongly Correlated Pupil Responses for all 30 Spots

To examine responses to the video messages, we first extracted the pupillary trace data for each video (in each condition) from every participant’s raw data file. Next, we combined all pupillary data for a given video/condition across all participants. Thus, for the first video (a commercial by Airbnb), we obtained 20 pupillary traces for the 100% video condition, 20 traces for the 50%-nd (no distraction)-condition, and 20 traces for the 50%-wd (with distraction) condition (yielding a total of 60 participants exposed to all variants of the Airbnb spot - and analogously for all other spots). We then plotted these data and computed the degree of inter-subjective response similarity. As can be seen from Figure 2, viewing the same video spot evokes fluctuations in pupil size that are shared among viewers, supporting our prediction.

**Figure 2:**
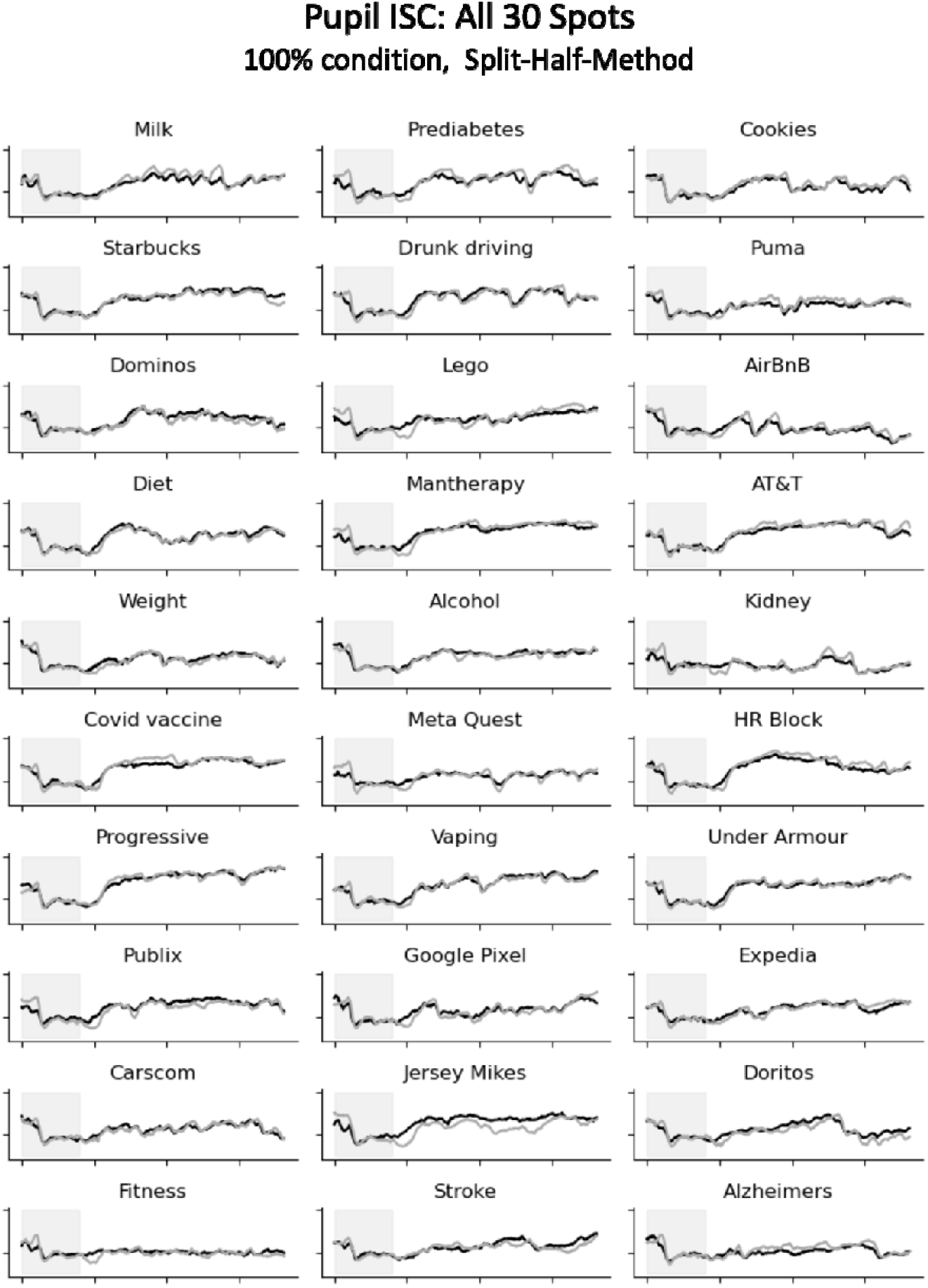
Pupillary response trajectories for all 30 spots (100% viewing condition). Black and gray time series show the average pupillary time series averaged across each sub-audience (n=10 viewers per line). As can be seen, the pupillary time series for each spot closely resemble each other across viewers exposed to the same spot, but they differ between spots. Gray lines mark the countdown/onset period of each video, which was excluded from the analysis.

Statistical analysis of these results confirmed highly significant pupil-ISC^3^ for 29 out of 30 spots. On average, the split-half correlation between pupil time series (in the fullscreen condition) was *r =* 0.78, with a maximum of *r =* 0.93 (for ‘kidney’) and a minimum of *r =* 0.08 (for the ‘fitness’-spot)^4^.

### Reception Data Enable Decoding of Spot Identity and Reveal the Influence of Viewing Condition

Having demonstrated that viewers’ pupils respond similarly to each video, we next examined the role of viewing condition. Specifically, we carried out the same analysis as for the 100%-viewing condition (see Figure 2) and also for the 25nd-and 25wd-conditions. As shown in Figure 3, this analysis revealed that even despite the different sizes (100 vs. 25%) and the varying perceptual demands (with vs. without distractors), the pupil traces still respond very reliably - and with a unique signature that is specific to each spot. Although Figure 3 only displays a subset of three exemplary spots (Milk, Prediabetes, and Cookies) due to space limitations, the pattern of results held again for the remainder (average correlation for 25nd *r = 0*.*74* and for 25-wd *r = 0*.*68*).

**Figure 3.**
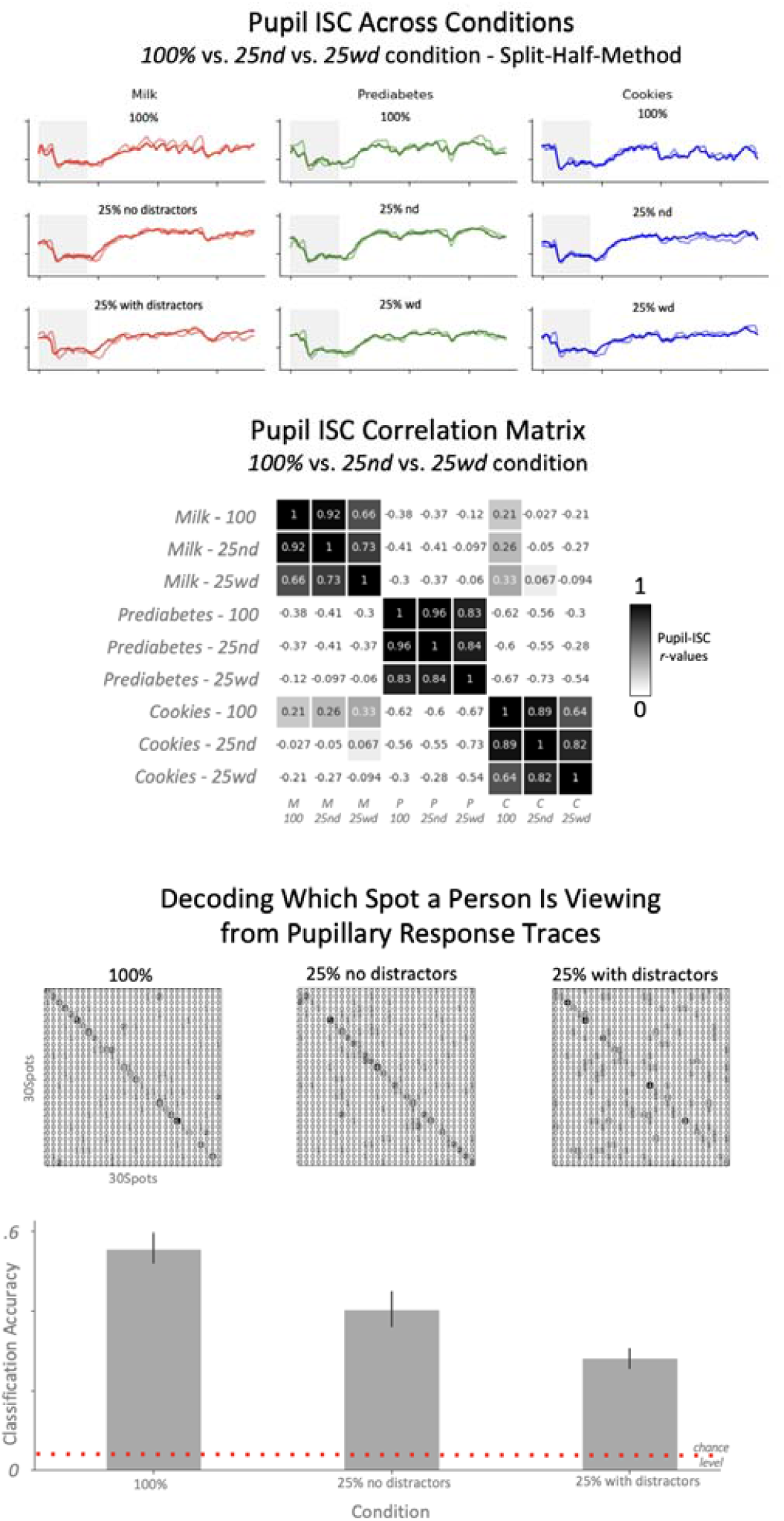
Pupil-ISC across viewing conditions and decoding accuracy. Top panel: Pupil-ISC results for three exemplary spots across conditions. Middle panel: When comparing the pupil-time-series from each condition for the three different spots, a robust block structure emerges along the diagonal. Thus, the same spot evokes a similar pupil signature over time, but different spots evoke a different signature. Bottom panel: Decoding analyses for all 30 spots. Given that pupil-time-series for a given spot are very similar to each other, it is possible to decode which spot a given individual is viewing from their pupil-time-series-data. The smaller confusion matrices exhibit a clear diagonal structure, and the bar graph at the bottom summarizes the overall decoding performance for each condition.

Critically, not only were the pupillary signatures for a particular spot correlated within each condition (e.g., across viewers exposed to the milk spot in the 25-wd-condition), but a given spot’s pupil response signature was also preserved across conditions: As shown in the middle panel of Figure 3, robust correlations of pupillary traces emerge for each spot (shown under different conditions), but correlations across the different spots vary, leading to the block-like diagonal structure.

Finally, we turned to the decoding question, which asked: Can we decode which spot a given individual is viewing based on their pupillary trace data? To answer this, we proceeded as follows: We combined all pupillary data (from a given condition) into a large dataset, one row for every pupillary time series, and labeled every row based on the video’s name. Then we applied machine learning methods (from the scikit-learn library and its time-series sublibrary sktime, specifically using a TimeSeriesForestClassifier with a 5-fold cross-validation) to fit a predictive model.

The results are illustrated in the bottom panel in Figure 3. We find that once the model is fit with pupillary time series data from other viewers, we can decode with relatively high accuracy which of the 30 spots an individual is viewing. Given that there are 30 spots, the chance level for this analysis would be 1/30 = 0.03. The achieved performance varied between 56% for the 100% condition), 40% for the 25%-nd condition, and 28% for the 25%-wd condition, and thus consistently above chance.

### Analysis of Memory Performance and Pupil-Memory Relationship

Having established that pupillary response time series are correlated across viewers and can b used to identify which spot a person is viewing, we next asked whether, beyond such general on audience-wide pupil responses, the pupillary data would also carry information about individual characteristics, particularly subsequent memory. To this end, we first compared overall per-spot memory (recall and recognition performance for a given spot - irrespective of pupil data). The results of this analysis are displayed in Figure 4.

**Figure 4.**
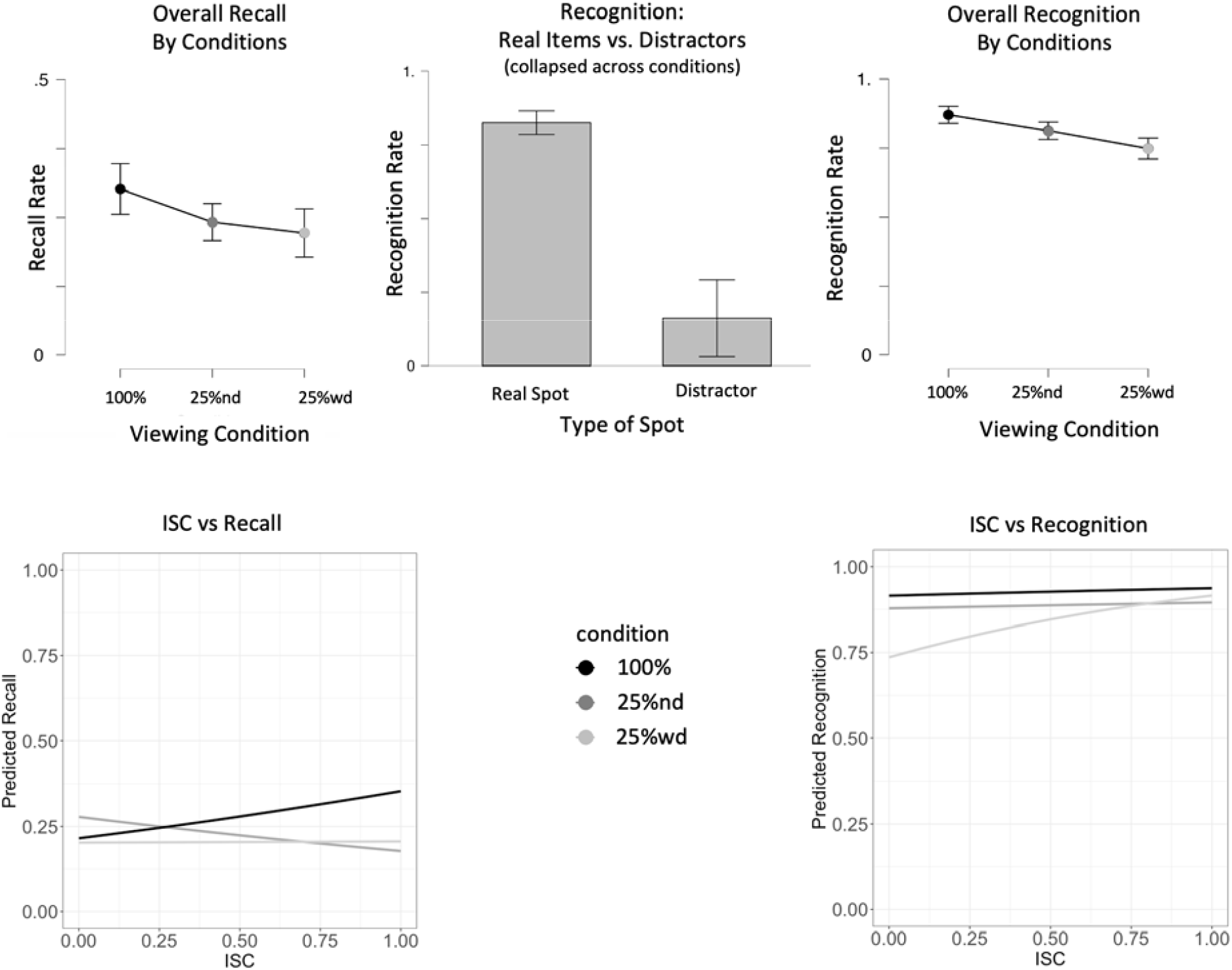
Memory Performance Across Viewing Conditions. Left panel: For the free recall, the condition in which a person was exposed to the spot (100%, 25%nd, 25%wd) strongly influenced recall. Of note, spots were assigned to different conditions for different participants hence this effect is not dependent on a specific spot’s memorability but is driven by the different presentation sizes (100% vs. 25%) or modes (with vs. without distraction). Central panel: For the recognition method, a few distractors were included to ensure that high recognition performance was not due to guessing. The marked difference between these distractors and the real memory items shows that guessing cannot explain the pattern of results. Right panel: The pattern of results for spot recognition matches the pattern for free recall. Bottom panels: Predicted effect plots from statistical analyses (see text for details).

For the free recall, we find that if a given spot was shown in the 100% condition, more participants recall it than when the same spot was shown in the 25%-nd or 25%-wd condition.^5^ A repeated-measures ANOVA (by-spots) revealed a significant main effect of *viewing condition, F = 4*.*2, p =* .*02*. On average, spots that were seen in the 100% condition were recalled by an average of *m*_*recall-100*_ = 6 participants (out of a maximum of 20). This was significantly higher compared to an average total recall of *m*_*recall-25nd*_ = 4.8 and *m*_*recall-25wd*_ *4*.*4* in the 25% conditions with or without distractors, respectively (see Figure 4).

A closely matching pattern of results was found when assessing memory via the recognition method. First, as expected, we find a far superior memory for spots that were actually presented compared to distractors (*mean recognition_rate*_*real_spots*_ *=* .*83, s*.*d. =* .*11; mean recognition rate*_*distractors*_ *=* .*16, s*.*d. =* .*18; t = 13*.*99, p <* .*001;* Figure 4, top middle panel; of note, we also found that the four thematically related distractors were more often falsely recognized than unrelated distractors, with average rates of *m*_*related*_ *=* .*23 vs. m*_*unrelated*_ *=* .*09*, respectively). Further exploration of the recognition rate based on the viewing condition in which a participant had been exposed to the spot again revealed a pattern that corresponds with the recall data: As shown in Figure 4 (top left panel), spots that were viewed in the 100% condition were recognized best, followed by spots in the 25%nd condition, and finally the 25%wd condition. Compared to the recall, which was relatively rare (∼15-35%), recognition performance was much higher (between ∼70-90%). Critically, the difference in performance based on viewing condition was highly significant (*F*_*viewing_condition*_ *= 13*.*9, p <* .*001*).

Having established that the different viewing conditions led to varying levels of recall/recognition, we can already assume that pupillary responses and memory performance will be related. For instance, the bottom panel in Figure 3 demonstrates that spot-wise decodability is highest for spots viewed in the 100% condition, followed by the 25%nd and 25%wd conditions; this corresponds with the pattern of recall and recognition rates across conditions seen in Figure 4. However, aside from this general relationship between pupil response (decodability) and subsequent memory, we were interested in examining the pupil-to-memory relationship at the level of individual trials. More specifically, we computed an ISC value for every trial that indicated how similar this participant’s pupillary signature was to the rest of the group (i.e., to the pupillary trace of all other participants who viewed the same spot under the same condition). The underlying reasoning is that the degree of inter-subjective similarity might serve as an index of neurotypical processing of a given spot. We then combined these trial-level ISC results with the information about memory, i.e., whether the spot was recalled/recognized or forgotten by this given individual, along with information about the condition, the spot, and the specific participant. The logic of this analysis is based on prior work that showed that when brain regions involved in memory encoding respond more reliably across viewers of specific TV scenes, then participants will be more likely to recognize that scene (Hasson et al., 2008). To examine this, we specified a generalized linear mixed-effects regression model using recognition (or recall) as a binary outcome (Bates et al., 2015), testing for the impact of ISC on recognition (or recall) probability while accommodating for influences of condition (modeled as a fixed effect) as well as spot-, and subject-specific differences (modeled as random effects). These analyses revealed a mixed pattern of results: At a global level, higher ISC was generally associated with higher recognition (or recall), as can be expected from the fairly large between-condition ISC differences and the parallel differences in spot recognition (or recall). However, consideration of the differences between conditions, spots, and subjects suggested a number of interactions, such as a stronger relationship between ISC and recognition for the 50wd condition (see Figure 4). Specifically, for the recognition outcome, we observed a marginally significant interaction (*p* = 0.07) between ISC and condition on recognition performance, along with a highly significant main effect of condition (due to the overall higher recognition results in 100 vs. 25%nd vs. 25%wd, *p* < 0.001). With regard to the fixed effects, we found that the relationship between ISC and recognition was greater in the 25%wd condition compared to the 25%nd and 100% condition. For the recall outcome, the pattern was generally similar, with a significant interaction between ISC and condition, *p* = 0.03, yet without a significant main effect of condition, *p* = 0.23. However, we note that the level of recall was generally low, which leads to fewer positive occurrences. Overall, these results suggest that the condition manipulation is associated with ISC, and pupillary ISC for given spots can be linked to memory outcomes. However, more complex interactions involving the level of ISC, the condition, and the memory outcome suggest that more work is needed to examine how pupil-ISC interacts with task-, subject-, and stimulus-level factors.

## Discussion

In this study, we exposed participants to media messages while capturing their pupillary responses. Our main goal was to study the inter-subject similarity of the pupillary traces for each video and to test whether it is possible to decode from the pupillometric response which message a given receiver/viewer is exposed to.

The results of this study are clear and can be summarized as follows: The pupillometry data are very robustly correlated across recipients of the same video. The degree of similarity (ISC) is affected by the presentation condition, but it is still possible to decode with high accuracy which video a person is viewing based on their pupillary process data, even across different exposure conditions. Finally, we examined the relationship between pupillary responses and subsequent memory, finding some evidence that pupillary ISC is related to subsequent memory. Overall, these findings support the validity and potential of this novel approach.

Perhaps the most notable finding is that it is possible to decode which of the 30 videos a person is viewing by looking at how their pupil fluctuates during reception. This ability to decode a spot’s identity is a consequence of the fact that different viewers’ pupils respond with similar pupil-size fluctuations to the same content. Thus, by creating a sort of database of typical pupil responses to video content, we can create a characteristic profile of the pupillary response fluctuations. We refer to this profile as the “pupil pulse” or pupillary response signature of a given video. Much like a barcode, a fingerprint, or an iris scan are signatures of personal identity, this is a characteristic profile of message identity. Because different videos have different physical characteristics, it is possible to database pupil response signatures from other viewers and then compare incoming reception data to the stored templates to decide which video the current person is likely viewing. In this way, pupillometry offers a promising method to connect media content and reception response. All that is needed for this is information about when a video starts, which is technically very feasible to obtain in media measurement.

### Broader Implications: Pupillometry for Audience Response Measurement

Looking at the field of communication and media psychology from a bird’s-eye perspective, one can notice several theoretical fault lines and measurement gaps between content, reception, and effects analyses (Schmälzle & Huskey, 2023). Specifically, the concept of exposure - whether an individual message is received by a given individual - lies at the foundation of mass communication and media effects (Hornik, 2002). Put simply, if a message fails to meet the eye, it cannot have any effect. This simple causal logic applies to media influence in general as well as specific effects like those of commercial advertising. Accordingly, the field has amassed elaborate audience metrics, such as readership estimates for print, TV ratings for viewership, or social media analytics and digital trace data (Peng et al., 2017), and elaborate statistical methods strive to incorporate exposure data in moderation and linkage analyses (de Vreese & Neijens, 2016; Kranzler et al., 2019). Yet, one must acknowledge that these varied methods often suffer from key gaps in how they ascertain actual exposure, thus failing to unequivocally establish the causal chain (Spencer et al., 2005) from media content to reception mechanisms and on to media. For instance, aggregated TV viewership data leave it unclear whether a given individual actually ‘took in’ a given message or whether they were present in another room but had the TV running. Likewise, even though social media enables tracking of large-scale audience behaviors, it is far from clear whether these all reflect real peoples’ activities^6^. With this in mind, the ability to close these gaps and zoom in on the exposure-reception-retention nexus is the main theoretical contribution of the approach presented here: By tracking the pupil dilations of individual viewers, we can conclude that their eyes were following the continuous unfolding of the media stimulus, and we can relate this data to subsequent memory.

However, aside from these theoretical considerations, an improved ability to rigorously quantify exposure and reception responses is not only theoretically significant, but will likely have substantial practical significance: Specifically, this new approach appears promising for next-generation media measurement^7^, particularly the Metaverse, and we can already expect that advertisers in the new media ecosystem will have a large interest in harvesting such user data (much like what happened with ‘cookies’ on the internet). Obviously, this development has important ethical and privacy implications that require our attention.

### Strengths, Limitations, and Avenues for Future Research

The strengths of the current study relate mainly to its particular mix of innovative methodologies for measuring, analyzing, and interpreting media-evoked responses in a moment-to-moment and unobtrusive manner. We also view the use of VR as promising because it comes with on-board measurement potential and allows to perfectly standardize and potentially manipulate environmental conditions; however, the approach can also work outside of VR and with stationary or mobile eye-tracking.

Given the novelty of this approach, many potential limitations could be mentioned. For instance, we only tested short video messages, and while we used a decent sample (30 messages and 60 viewers), questions regarding generalizability remain. Likewise, our choice to present normal videos and two altered versions (one reduced to 25% size, the other reduced accompanied by distractors) represents just one of many options for manipulations of the stimulus, requiring many others to follow. For instance, one could ask if the results would be the same had we also turned the videos to grayscale, enlarged or shrunk them further, or added user-sided attention manipulations, and so forth. Clearly, although our results are positive and promising, there are also many open questions with regard to further psychological influences beyond the more stimulus-driven pupillary-light reflex. For instance, if we take the view of advertisers who might be interested in, e.g., targeting or tailoring of messages, which is a relevance-based manipulation, then one might ask how this affects pupillometric data in addition to, or in interaction with the PLR.

Despite these limitations, further exploration holds significant promise. Avenues that appear highly favorable include the integration of pupillometric measures with other data types, most notably eye coordinates, but also a broader set of neurocognitive data. Furthermore, we opted for the use of a TV viewing room to expose viewers under standardized conditions, but from a practical perspective, exposure would likely occur in varying circumstances, which could also be examined. Finally, the primary focus of this study was on the video (message identity), with a secondary focus on incidental memory for those videos. Now that the paradigm and robust results are established for these basic foci, future work can zoom in on more nuanced psychological reception processes in between (e.g., interest, engagement, and so forth).

### Summary and Conclusion

In conclusion, the eye is at the vanguard of the reception process, representing the place where visual information arrives and is transformed into neural signals. The Pupillary Light Reflex (PLR) is one of the first and most robust effects media have on the neurocognitive system. This study exposed audiences to visual content and used pupillometric traces to decipher which video individuals viewed. This approach could have important practical applications in media and advertising response measurement. Moreover, because the eye represents the actual point of contact between an external media stimulus and the subject-sided reception response, it is the place where “exposure” happens, and “reception” begins. Given the enormous significance of this exposure-reception nexus for all forms of mass-mediated communication, a principled ability to objectively and unobtrusively capture relevant data from viewers, and with an eye towards collectively shared audience responses, holds significant promise for tightening the theoretical link between media content and effects.

## Open practices, data, and code availability

The data and materials for this study are available at https://anonymous.4open.science/r/vr_video_pupil_study-0CEE/. Additional VR-related scripts will be made available to interested researchers.

## Footnotes

1 To be clear, we do not dispute that for many cognitive science applications, the physical stimulus characteristics that drive the PLR are, in fact, a confound. Thus, we do not dispute that it makes typically much sense to control for these characteristics to more purely assess cognitive modulations of pupil dilation. However, the core argument of this paper is that for some applications, particularly for audience measurement, the PLR is not necessarily a confound/noise, but can serve as a source of high-quality signal.

2 Across the different conditions and measures, data from some participants had to be excluded, although such cases were rare: For instance, for some pupil-ISC analyses, there should be 20 participants per viewing condition (100 vs. 25nd vs. 25wd, i.e., 3*20=60). However, because of technical failure, combined with the randomized presentation of spots, not all 30 pupil-spot data were available. This led to random missing data for a few spots (i.e., values can be based on 19 instead of 20 values). However, given the random presentation rate and the few instances in which such crashes occurred, the overall data loss was very low, and certainly far lower compared to other measures (e.g., ECG, EEG, or fMRI, where regularly 10-30% of data are missing).

3 Note that for this analysis, we rely on the split-half analysis method in which the group of viewers is divided int two halves and ISC is computed across the averages. ISC is also high and significant if assessed at the level of eac individual, as will become apparent in the analyses below (specifically the predictive model). However, computin ISC across groups helps overcome individual measurement noise and allows to better demonstrate the results. Mathematically, the procedures are related and the group ISC is generally higher, but ranking is preserved.

4 Of note, we computed these analyses after excluding the onset-transient that is marked by the shaded regions; these onset-transients comprised the 3-2-1-countdown video, the black transition screen, as well as the first seconds of the actual spot.

5 Note that because different participants viewed different spots in different conditions, this analysis controls for the varying memorability of certain spots.

6 For instance, in the media measurement industry, it is very common to mark media content in order to identify how often people consume specific content. Unique landing pages and web cookies are good examples of these widespread practices. For streaming content, these can be equipped with specific markers as well. However, this practice is laborious, costly, and the flood of ever-changing online content like ads, and short videos is an obstacle to these practices though. With this in mind, the approach offers a fresh way of thinking about this problem: Instead of marking the content, one can compare how similarly different people respond to the same content. That way, content does not have to be marked, but it is possible to decode what people are viewing; and an added benefit is that such receiver-sided bio- and neurometric data could predict outcomes like memory, especially when used at scale.

7 but it clearly also applies to traditional media formats.

